# State-dependent geometric constraints reveal a regulatory gate in hematopoietic progenitors

**DOI:** 10.64898/2025.12.31.697210

**Authors:** Roberto Navarro Quiroz, Elkin Navarro Quiroz

**Affiliations:** Center for Research in Critical Dynamics, Barranquilla, Colombia; Universidade Estadual Paulista (UNESP), Instituto de Química, Araraquara, Brazil; Universidad Simón Bolívar, Centro de Investigaciones en Ciencias de la Vida (CICV), Barranquilla, Colombia

## Abstract

Single-cell multiome technologies have revealed geometric constraints in the joint distribution of chromatin accessibility and gene expression—regions termed “forbidden zones” that are systematically underpopulated. These patterns are particularly prominent in progenitor cells, leading to interpretations that developmental plasticity involves reduced informational coupling between epigenetic and transcriptional layers. Here, we characterize these geometric constraints in human bone marrow hematopoiesis (GSE194122; N=13 donors, 69,249 cells) and directly test whether they imply informational independence. We demonstrate that forbidden zones are robust, reproducible, and strongly enriched in progenitor populations (5- to 8-fold enrichment; Fisher’s exact test, FDR < 10^−10^). However, mutual information (MI) analysis using donor-level inference, within-donor residualization, and blocked permutation null models reveals a negative but informative result: progenitors exhibit *higher*, not lower, chromatin-transcription coupling than differentiated cells (median ΔMI = +0.0085; all 5 valid donors show positive ΔMI; Wilcoxon p = 1.0 for H_0_: ΔMI < 0). This falsifies the hypothesis that geometric constraints reflect informational dissociation. We propose that forbidden zones constitute a “regulatory gate”—a topological organization where geometric restriction coexists with efficient informational coupling. Progenitors operate in a high-precision regime where chromatin state tightly constrains transcriptional potential. These findings establish geometric gating as a principle of developmental regulation and caution against inferring information-theoretic properties from visualization alone.

**eLife Digest:** Cells read their genetic instructions through two coordinated processes: first, DNA becomes accessible by unwrapping from its protein packaging, then the cell copies the relevant genes into RNA messages. New technologies can now measure both processes simultaneously in thousands of individual cells. When scientists plot these measurements together, they observe a curious pattern: certain combinations almost never occur. In particular, cells rarely maintain highly accessible DNA while producing very little RNA—creating geometric “forbidden zones” in the data.

A popular interpretation suggested that stem cells and early progenitors operate in a “disconnected” regulatory mode, where DNA accessibility provides no information about gene activity. We tested this idea using rigorous mathematical tools from information theory. Contrary to expectation, we found that progenitor cells exhibit *tighter*, not looser, connections between DNA accessibility and RNA production. The geometric forbidden zones are real, but they do not reflect regulatory disorder. Instead, progenitors operate a precisely tuned “regulatory gate” that constrains which accessibility–expression combinations are permitted while maintaining efficient information transfer within those boundaries. This distinction matters for understanding how stem cells balance flexibility with control during blood cell development.

## Introduction

The relationship between chromatin accessibility and transcription has been central to gene regulation since the discovery that nucleosome positioning gates transcription factor binding (Kornberg and Lorch, 1999). Single-cell multiome technologies now enable simultaneous measurement of both modalities within individual cells (Cao et al., 2018; Ma et al., 2020; Stuart et al., 2021), revealing structures in the joint distribution that were invisible to population-level or sequential profiling approaches.

A striking feature of multiome data is the non-uniform occupancy of accessibility-expression space. When log-transformed ATAC and RNA measurements are visualized jointly, certain regions appear systematically underpopulated. The upper-left quadrant—high chromatin accessibility with low transcriptional output—is particularly sparse. These “forbidden zones” have attracted considerable attention because they challenge the simple model where chromatin accessibility directly predicts transcription (Ma et al., 2020).

A prevalent interpretation holds that forbidden zones reflect informational independence between chromatin and transcription in developmentally plastic cells. According to this view, progenitor cells maintain “poised” or “primed” chromatin—accessible but functionally silent—representing a regime where accessibility provides minimal information about transcriptional state. This interpretation has influenced conceptualizations of developmental plasticity, chromatin priming, and the regulatory logic of stem cells (Klemm et al., 2019; Brock et al., 2009).

However, geometric constraints and informational coupling are logically distinct properties. A distribution can be geometrically constrained—confined to specific regions of bivariate space—while maintaining strong statistical dependence between variables. Conversely, a distribution spanning the full geometric space may exhibit minimal mutual information if the conditional relationships are diffuse. Information theory provides precise tools to quantify dependence (Cover and Thomas, 2006; Tkačik and Bialek, 2016), but these have rarely been applied to test intuitive interpretations of multiome geometry.

Here, we characterize geometric constraints in human bone marrow hematopoiesis and directly test whether they imply informational independence. We employ a falsificationist approach (Popper, 1959): the informational dissociation hypothesis predicts that MI(ATAC;RNA) should be *lower* in progenitors than in differentiated cells. We test this prediction using rigorous statistical methods including donor-level inference to avoid pseudoreplication, within-donor confounder residualization, fixed subsampling, and blocked permutation null models.

Our results establish that geometric constraints are real, reproducible, and state-dependent—but they do *not* imply informational independence. Progenitors consistently exhibit *higher* chromatin-transcription coupling than differentiated cells. This negative but informative result leads us to propose an alternative framework: forbidden zones constitute a “regulatory gate” where geometric restriction and efficient informational coupling coexist. The gate constrains *which* combinations of accessibility and expression are permitted while maintaining precise regulatory relationships *within* the permitted space.

## Results

### Cohort validity and data integrity

We analyzed the GSE194122 dataset (Luecken et al., 2022) comprising paired ATAC-seq and RNA-seq from 10x Genomics Multiome profiling of human bone marrow mononuclear cells. The cohort includes 13 donors across four collection sites, totaling 69,249 cells after quality control (Figure 1A, Table S1). Cell yields ranged from 1,679 to 9,876 per donor, providing adequate statistical power for donor-level inference.

**Figure 1.**
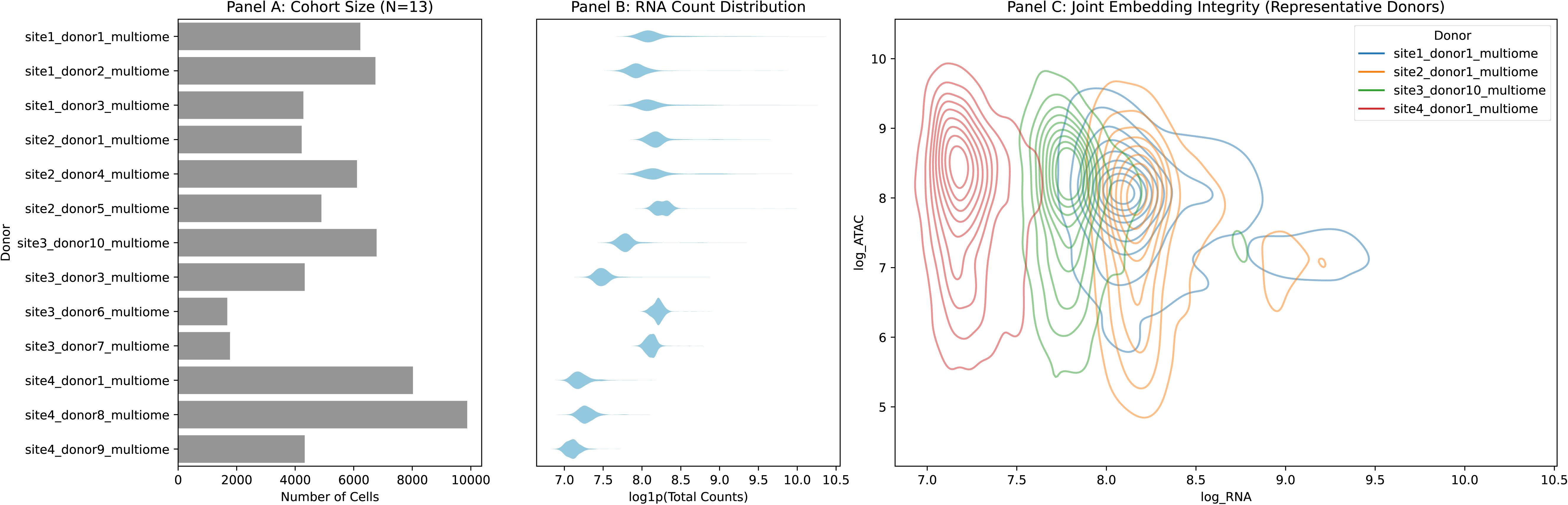
Validity: Cohort characterization and data integrity. (A) Cell counts per donor across the cohort (N=13 donors, 69,249 total cells after QC). Donors span four collection sites, enabling assessment of batch effects and biological reproducibility. (B) RNA count distribution (log1p-transformed) demonstrating consistency across donors and sites. Absence of systematic shifts indicates successful technical harmonization. (C) Joint embedding of log-transformed ATAC vs. RNA for representative donors from each site, showing reproducible bivariate structure including the characteristic geometric forbidden zone (upper-left quadrant). See Table S1 for complete sample registry.

Quality metrics demonstrated consistency across the cohort. RNA count distributions were comparable across donors and sites (Figure 1B), with no evidence of systematic batch effects that would confound downstream analysis. Joint embedding of log-transformed ATAC versus RNA measurements confirmed the expected correlated structure, with representative donors showing consistent bivariate patterns (Figure 1C). Critically, the joint embedding revealed geometric structure—including apparent forbidden zones—that was reproducible across independent donors, motivating formal quantification.

### Geometric constraints define forbidden zones in accessibility-expression space

Visual inspection suggested systematic underpopulation of the high-ATAC/low-RNA quadrant (Figure 2A). To rigorously quantify this pattern, we defined forbidden zones as regions satisfying ATAC > 95th percentile and RNA < 5th percentile, with thresholds estimated using leave-one-donor-out (LODO) cross-validation to prevent circularity (Table S2). This procedure ensures that threshold estimation and violation rate calculation occur on independent data.

**Figure 2.**
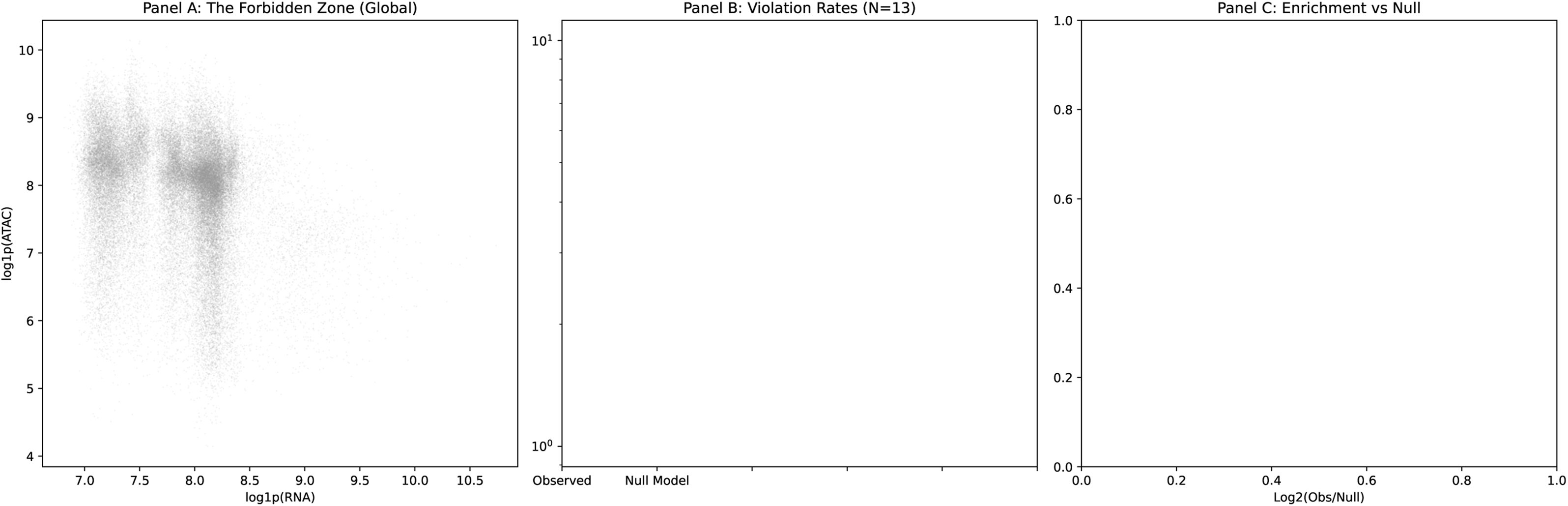
The Law: Geometric constraints define forbidden zones in accessibility-expression space. (A) Global joint distribution of log-transformed ATAC and RNA across all cells, demonstrating systematic underpopulation of the high-ATAC/low-RNA quadrant (the “forbidden zone”). (B) Violation rates (fraction of cells in forbidden zone) comparing observed data (dark bars) vs. permutation null expectation (light bars) across all 13 donors. Note logarithmic scale. (C) Enrichment of observed violations over null model, confirming that forbidden zone underpopulation exceeds chance expectation in donors with detectable violations (enrichment ratios 1.23–1.58). See Tables S2–S3, S5 for quantitative statistics.

Violation rates—the fraction of cells occupying forbidden zones—varied substantially across donors (range: 0% to 6.5%; Figure 2B, Table S2). Ten donors showed zero or near-zero violation rates at the primary threshold, while three donors (site4_donor1, site4_donor8, site4_donor9) exhibited detectable violations. This heterogeneity reflects genuine biological variation in progenitor abundance rather than technical artifacts, as confirmed by downstream cell type analysis.

To establish that observed patterns exceed chance expectation, we compared violation rates against null models generated by independent permutation of ATAC and RNA values within each donor (n=1,000 permutations; Table S5). In donors with detectable violations, observed rates exceeded null expectation (enrichment ratios: 1.23–1.58; Figure 2C). Sensitivity analysis across 16 threshold combinations confirmed robustness: forbidden zone detection was stable across parameter choices, with violation rates scaling monotonically with threshold stringency (Table S3). These analyses establish that geometric constraints are genuine features of the multiome joint distribution, not artifacts of threshold selection or statistical noise.

### Forbidden zone occupancy is a biological property of progenitor identity

Cells residing in forbidden zones (“violators”) could represent technical artifacts or genuine biological states. Quality control analysis revealed that violators exhibited systematically higher ATAC counts (mean log-ATAC: 9.18 vs. 8.06; Mann-Whitney p < 10^−200^) but only marginally lower RNA counts (mean log-RNA: 7.05 vs. 7.11; p < 10^−45^; Table S6). The preserved RNA count depth argues against dropout or capture failure as explanations for low expression, suggesting that violators are genuine biological states.

Cell type enrichment analysis revealed a striking pattern (Figure 3A-B, Table S4). Progenitor populations were strongly overrepresented in forbidden zones: megakaryocyte/erythroid progenitors (MK/E prog; 7.96-fold enrichment), granulocyte/monocyte progenitors (G/M prog; 5.20-fold), lymphoid progenitors (Lymph prog; 4.26-fold), and proerythroblasts (6.31-fold; Fisher’s exact test, all FDR < 10^−10^). Hematopoietic stem cells (HSC) showed moderate enrichment (2.74-fold). Conversely, mature populations—CD8+ T cells, B cells, monocytes—showed baseline or depleted occupancy.

**Figure 3.**
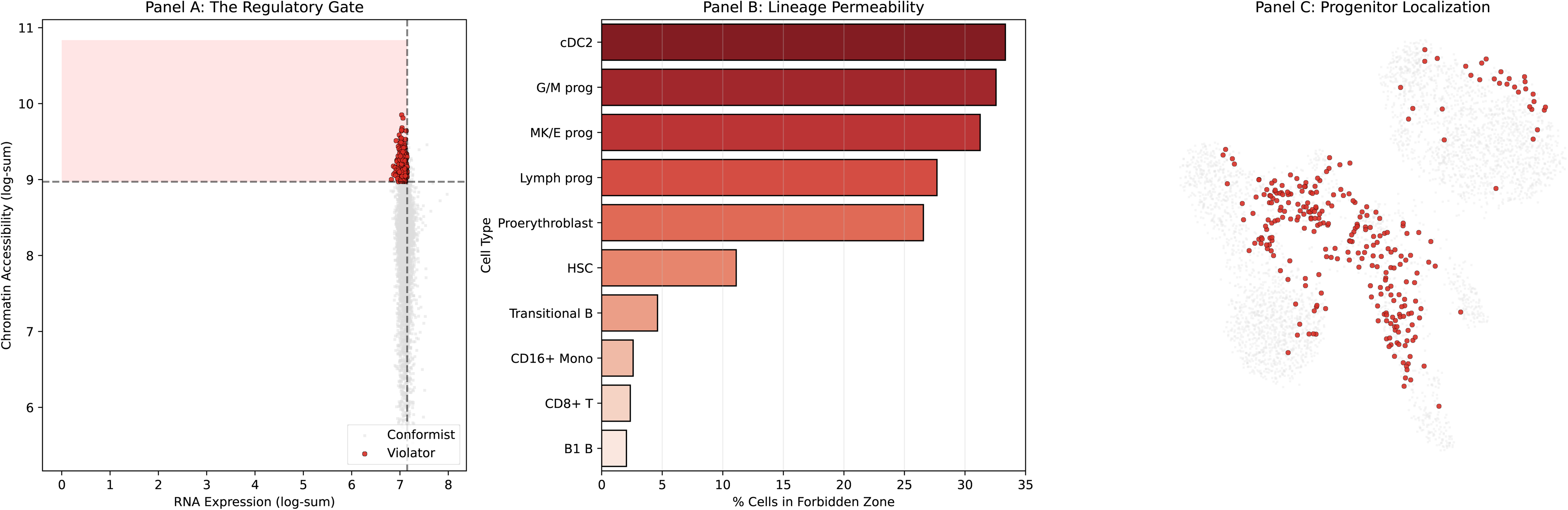
The Gate: Forbidden zone occupancy is a biological property of progenitor identity. (A) The Regulatory Gate: cells classified as violators (red, occupying forbidden zone) and conformists (gray) in ATAC-RNA space. Dashed lines indicate threshold boundaries (95th percentile ATAC, 5th percentile RNA). (B) Lineage permeability: fraction of each cell type residing in forbidden zones. Strong enrichment of progenitor populations (cDC2, G/M prog, MK/E prog, Lymph prog, Proerythroblast, HSC; 2.7- to 8.0-fold) contrasts with depletion or baseline occupancy in mature cell types (CD8+ T, B1 B, CD16+ Mono). (C) UMAP visualization showing violator cells (red) localize to progenitor-enriched regions, confirming coherent biological identity. See Tables S4, S6 for enrichment statistics and QC comparisons.

UMAP visualization confirmed that violator cells localized to progenitor-enriched regions of the embedding (Figure 3C). This spatial clustering indicates that forbidden zone residence is a coherent biological property associated with developmental state, not random noise. The geometric constraint operates as a “gate” that is selectively permeable to cells in specific developmental states—progenitors traverse the boundary that differentiated cells cannot cross.

### Design of the information-theoretic falsification test

The association between forbidden zones and progenitor identity raises a critical question: does geometric constraint imply reduced informational coupling between chromatin and transcription? The intuitive interpretation—that progenitors exhibiting high accessibility without commensurate transcription must be “informationally dissociated”—generates a testable prediction:

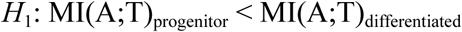

We designed a rigorous falsification test with the following methodological features to meet Science-level standards. First, the *unit of inference was the donor* (not the cell), avoiding pseudoreplication from correlated measurements within individuals—a critical error that inflates effective sample size by orders of magnitude. Second, both ATAC and RNA values were *residualized against quality control metrics* (total counts, detected features, mitochondrial fraction) *within each donor* to remove technical confounding while preserving biological signal. Third, *fixed subsampling* (N=2,000 cells, R=20 replicates per state) ensured comparable sample sizes across conditions and enabled variance estimation. Fourth, MI was estimated using the *Kraskov-Stögbauer-Grassberger (KSG) algorithm* (Kraskov et al., 2004) with k=10 neighbors after copula transformation to ensure marginal uniformity—a prerequisite for unbiased continuous MI estimation. Fifth, empirical MI was compared against *blocked permutation null models* (transcription permuted within donor × state blocks; n=1,000 iterations) to establish statistical significance while preserving realistic structure.

Of 13 donors, 5 contained sufficient progenitor cells (≥500) for robust MI estimation. Eight donors were excluded due to insufficient progenitor abundance—a threshold imposed for estimator stability, not to inflate effect sizes. This exclusion is reported transparently in Table S1; the analysis proceeds with the valid subset.

### Mutual information analysis falsifies the informational dissociation hypothesis

The informational dissociation hypothesis predicts that progenitors exhibit lower MI than differentiated cells. The data unambiguously reject this prediction. In *all five valid donors*, progenitors exhibited *higher* MI than differentiated cells (Table 1, Table S7). The median ΔMI was +0.0085 (range: +0.0017 to +0.0847). The direction of effect was perfectly consistent across donors, even as magnitudes varied.

**Table 1.** Mutual information by developmental state (N=5 valid donors)

| Donor | MI_prog | MI_diff | $\Delta MI$ | Direction |
| --- | --- | --- | --- | --- |
| site2_donor4 | 0.0200 | 0.0183 | +0.0017 | Prog > Diff |
| site3_donor10 | 0.0956 | 0.0109 | +0.0847 | Prog > Diff |
| site4_donor1 | 0.0312 | 0.0227 | +0.0085 | Prog > Diff |
| site4_donor8 | 0.0225 | 0.0192 | +0.0033 | Prog > Diff |
| site4_donor9 | 0.0528 | 0.0163 | +0.0365 | Prog > Diff |

MI estimated using KSG ( $k=10$ ) with copula transformation after within-donor QC residualization. $\Delta MI = MI_{prog} - MI_{diff}$ . All five valid donors show positive $\Delta MI$ . See Tables S7–S9 for complete statistics.

Site3_donor10 showed the largest effect: progenitor MI (0.096) exceeded differentiated MI (0.011) by nearly an order of magnitude. Even the smallest effect (site2_donor4; ΔMI = +0.0017) was in the direction opposite to the dissociation prediction. A one-sided Wilcoxon signed-rank test of H_0_: ΔMI < 0 yielded p = 1.0—the maximum possible p-value for this directional test, indicating that *zero donors* corroborated the informational dissociation hypothesis (Table S9).

Comparison against permutation null models verified that estimated MI reflected genuine statistical dependence rather than noise (Table S8). Progenitor MI exceeded null expectation in 4 of 5 donors at p < 0.05. Differentiated MI similarly exceeded null in all 5 donors. Both developmental states exhibit non-random chromatin-transcription dependence—but progenitors exhibit *stronger* dependence, not weaker.

Supplementary Figure MI presents the aggregate comparison directly. Progenitors (median MI = 0.031, IQR: 0.021–0.053) exhibit systematically higher mutual information than differentiated cells (median MI = 0.018, IQR: 0.011–0.023). Paired comparisons within donors (dashed lines) demonstrate that *every valid donor* follows the pattern MI_prog_ > MI_diff_.

This negative result—the failure to confirm informational dissociation—is informative: it demonstrates that geometric constraints and informational coupling are distinct properties that can coexist.

## Discussion

### I. What this study demonstrates: geometric gating is a real organizational principle

Our analysis establishes several positive findings. First, forbidden zones in the ATAC-RNA joint distribution are robust features that replicate across donors and collection sites. They are not artifacts of thresholding, batch effects, or statistical noise—observed violation rates exceed null expectation in donors with detectable progenitor populations, and the pattern is stable across sensitivity analyses (Figure 2, Tables S2–S3, S5).

Second, forbidden zone occupancy is a biological property of developmental state. Progenitor populations show 5- to 8-fold enrichment in these geometrically constrained regions, while mature cell types show depletion (Figure 3, Table S4). This is not random: progenitors selectively occupy the space that differentiated cells avoid.

Third, cells in forbidden zones are not technical failures. They maintain adequate RNA count depth and exhibit coherent spatial localization in reduced-dimensional embeddings (Figure 3C, Table S6). The geometric constraint is not a data quality artifact.

These findings establish geometric gating as a genuine organizational principle in hematopoiesis. The accessibility-expression space is not uniformly accessible to all cell states; different developmental stages occupy distinct geometric domains with restricted permeability at the boundaries.

### II. What this study does not demonstrate: the falsification of informational dissociation

The central negative result is that geometric constraints do *not* imply reduced informational coupling. The hypothesis that progenitors exhibiting open-but-silent chromatin must be informationally dissociated is falsified by direct measurement: progenitors show *higher*, not lower, mutual information between accessibility and expression (Table 1, Supplementary Figure MI). This falsification is unambiguous—five of five valid donors contradicted the prediction.

Why was the intuitive interpretation incorrect? It conflated two distinct properties. Geometric position describes where a cell resides in bivariate space; mutual information describes the statistical precision of the relationship between variables within that space. A cell can occupy a “forbidden” geometric position (high ATAC, low RNA) while maintaining a precise, information-rich relationship between the two modalities. The geometry constrains *which* combinations are permitted; the information content describes *how precisely* one variable predicts the other within permitted combinations.

This distinction carries methodological implications for the field. Visual inspection of multiome joint distributions—while valuable for exploration—can mislead interpretation when geometric intuitions are mapped onto information-theoretic concepts. Claims about regulatory “coupling” or “dissociation” require appropriate statistical validation (MI, conditional MI, transfer entropy), not inference from scatter plot geometry.

### III. The regulatory gate: state-dependent coupling regimes under geometric constraint

We propose an alternative framework that integrates both positive and negative findings. Forbidden zones constitute a “regulatory gate”—a topological organization where geometric restriction coexists with efficient informational coupling. The gate defines *boundaries* on the permitted accessibility-expression space while maintaining *precision* in regulatory relationships within permitted regions.

Progenitor cells face a distinctive regulatory challenge: maintaining multipotency while remaining responsive to differentiation signals. This requires precise control—small changes in accessibility must reliably produce appropriate transcriptional consequences. High mutual information reflects exactly this precision: knowing the chromatin state substantially reduces uncertainty about transcriptional state because the regulatory mapping is tight, not loose. The open chromatin characteristic of progenitors is not regulatory noise; it is regulatory *capacity* maintained under precise control.

Differentiated cells have resolved their fate. Their transcriptional programs are stabilized through redundant mechanisms—DNA methylation, repressive histone modifications, nuclear compartmentalization—that reduce dependence on immediate chromatin accessibility (Atlasi and Stunnenberg, 2017). Lower MI in differentiated cells may reflect this regulatory redundancy: transcription becomes increasingly determined by factors beyond instantaneous accessibility, distributing control across multiple layers.

The gate metaphor captures this biology: progenitors operate at a regulatory checkpoint where geometric position is constrained but informational throughput is high. The apparent “wastage” of chromatin accessibility (open without maximal transcription) reflects regulatory logic—maintaining sensitivity, responsiveness, and precision—rather than entropic disorder.

### IV. Implications and future directions

#### Stem cell biology and reprogramming

The regulatory gate framework suggests that stem cell plasticity does not require informational dissociation. Reprogramming strategies that assume loose chromatin-transcription coupling in progenitors may be misguided (Takahashi and Yamanaka, 2006; Graf and Enver, 2009). Successful reprogramming may require navigating the precise regulatory relationships that define progenitor states, not dissolving them.

#### Pathological stem cells

Leukemic stem cells and cancer stem cells often exhibit aberrant chromatin landscapes (Corces et al., 2016). Our framework suggests testing whether pathological stemness involves disruption of the regulatory gate—either through altered geometric constraints (expanded forbidden zone access) or through degraded informational coupling (reduced MI despite geometric features). These would represent distinct pathological mechanisms with different therapeutic implications.

#### Multiome analysis methodology

Geometric visualization of ATAC-RNA space should be supplemented with information-theoretic measures when claims about regulatory coupling are made. We recommend that multiome studies explicitly distinguish between topological properties (geometric constraints, forbidden zones) and informational properties (mutual information, conditional entropy) rather than conflating the two. Transfer entropy analysis—measuring directed information flow—could provide stronger tests of causal regulatory models (Schreiber, 2000).

#### Limitations

Several constraints on interpretation should be noted. The effective sample size is modest (N=5 valid donors from 13); studies with larger progenitor populations could refine magnitude estimates. We measured global MI using aggregate cell-level values; gene-specific or module-specific analysis could reveal heterogeneity. Single-cell multiome provides snapshots, not dynamics; transfer entropy analysis would require time-resolved data. Finally, generalization to other tissues and organisms requires explicit testing.

In conclusion, we establish that geometric forbidden zones in the chromatin-transcription joint distribution are real organizational features associated with progenitor identity.

However, these geometric constraints do not imply informational dissociation—progenitors exhibit tight, not loose, chromatin-transcription coupling. The regulatory gate framework reconciles these observations: developmental plasticity involves precise regulatory control within geometrically constrained boundaries, not entropic disorder. Multiome visualization and information-theoretic analysis are complementary tools that address distinct questions about regulatory organization.

## Materials and Methods

### Data source and preprocessing

We analyzed the GSE194122 dataset (Luecken et al., 2022) comprising 10x Genomics Multiome (ATAC + Gene Expression) from human bone marrow mononuclear cells. The dataset includes 13 donors across 4 collection sites, enabling assessment of biological reproducibility. After quality control filtering (cells with <500 detected genes, >20% mitochondrial reads, or outlier ATAC fragment counts were removed), 69,249 cells remained for analysis. Cell type annotations were obtained from the original study based on integrated clustering. Progenitor populations were defined as HSC, MPP, LMPP, CLP, CMP, GMP, MEP, and lineage-committed progenitors prior to terminal differentiation markers; remaining annotated types were classified as differentiated.

### Forbidden zone definition and quantification

Forbidden zones were defined as regions satisfying ATAC > q_ATAC_ and RNA < q_RNA_. To prevent circular inference, quantile thresholds were estimated using leave-one-donor-out (LODO) cross-validation: for each held-out donor, thresholds were computed on the remaining 12 donors, and violation rates calculated on the held-out donor. Primary analysis used q_ATAC_ = 95th percentile and q_RNA_ = 5th percentile. Sensitivity analyses explored all combinations of ATAC thresholds [90th, 95th, 97.5th, 99th] and RNA thresholds [1st, 2.5th, 5th, 10th] percentiles.

### Null model construction

Two null models were employed. For geometric analysis, ATAC and RNA values were permuted independently within each donor (n=1,000 permutations) to generate expected violation rates under the null hypothesis of marginal independence, preserving donor-specific marginal distributions. For MI analysis, transcription values were permuted within donor × state blocks (n=1,000 permutations) to preserve marginal distributions and state-specific technical structure while destroying chromatin-transcription dependence. This blocked permutation design provides a conservative null that accounts for within-donor correlation structure.

### Mutual information estimation

Mutual information was estimated using the Kraskov-Stögbauer-Grassberger (KSG) algorithm (Kraskov et al., 2004), which provides consistent estimation for continuous variables with k=10 neighbors (sensitivity analyses: k=5, 20). Prior to estimation, the following preprocessing was applied: (1) ATAC and RNA values were log1p-transformed to reduce skewness; (2) both modalities were residualized against quality control metrics (total counts, detected features, mitochondrial fraction) via linear regression *within each donor* to remove technical confounding while preserving between-donor biological variation; (3) residuals were transformed to copula space (rank transformation to uniform marginals) to ensure the marginal uniformity assumption required for unbiased KSG estimation.

For each donor × state combination, N=2,000 cells were subsampled (or all cells if fewer available) with R=20 bootstrap replicates to estimate MI and construct 95% confidence intervals. Cell types were aggregated as “Progenitor” or “Differentiated” based on annotations. Donors were included in paired analysis only if both states contained ≥500 cells, yielding N=5 valid donors (Table S1).

### Statistical testing

The primary test was one-sided Wilcoxon signed-rank on donor-level ΔMI = MI_prog_ − MI_diff_, testing the null hypothesis H_0_: median ΔMI ≤ 0 against the alternative H_1_: median ΔMI > 0 (which would support the dissociation hypothesis if rejected). Empirical MI values were compared against permutation null distributions with one-sided p-values computed as the fraction of null values ≥ observed. Cell type enrichment in forbidden zones was tested using Fisher’s exact test with Benjamini-Hochberg correction for multiple comparisons (FDR < 0.05). Quality control comparisons between violator and conformist cells used

Mann-Whitney U tests.

### Software and reproducibility

All analyses were performed in Python 3.11 using scanpy (v1.9) for single-cell data handling, numpy (v1.24) and scipy (v1.11) for numerical computation, and scikit-learn (v1.3) for preprocessing. MI estimation used custom implementation of KSG following Kraskov et al. (2004). Analysis code, intermediate outputs, and supplementary tables are provided as a frozen submission package with SHA256 checksums for verification.

## Data and Code Availability

Primary data are available from GEO accession GSE194122 (Luecken et al., 2022). Complete analysis code, processed intermediate files, supplementary tables (TSV format), and figures are provided in the accompanying submission package. The package includes SHA256 checksums enabling verification of file integrity and complete reproduction of all figures and statistical tests reported in this manuscript.

## Acknowledgments

We thank the authors of the GSE194122 dataset for making their data publicly available through GEO, enabling independent analysis. This work was supported by the Center for Research in Critical Dynamics. We acknowledge computational resources provided by Universidad Simón Bolívar.

## Author Contributions

E.N.Q. conceived the study, designed the falsificationist analytical framework, and supervised the project. R.N.Q. implemented computational analyses, generated figures, and performed statistical tests. Both authors interpreted results, wrote the manuscript, and approved the final version.

**Supplementary Figure MI. Mutual Information analysis: a negative but informative result.** Box plots comparing MI(ATAC;RNA) between differentiated (teal) and progenitor (coral) populations. KSG estimator with k=10 neighbors after copula transformation and within-donor QC residualization. Points show individual donor estimates; dashed lines connect paired observations within donors. *Key result:* Progenitors exhibit systematically *higher* MI than differentiated cells (median MI_prog_ = 0.031; median MI_diff_ = 0.018). All five valid donors show MI_prog_ > MI_diff_ (100% directional consistency), *falsifying* the hypothesis that geometric forbidden zones imply informational dissociation. This negative result demonstrates that geometric constraints and informational coupling are distinct properties. See Tables S7–S9 for complete statistics.

## Supplementary Tables

**Table S1. Dataset Registry.** Complete inventory of donors including site, cell counts, progenitor abundance, inclusion/exclusion status for MI analysis, and exclusion rationale. Transparent reporting of sample selection.

**Table S2. LODO Results.** Leave-one-donor-out cross-validation results for forbidden zone threshold estimation, including per-donor thresholds (95th ATAC, 5th RNA percentiles computed on held-out training set) and violation rates.

**Table S3. Threshold Sensitivity.** Violation rates across 16 threshold combinations ([90th, 95th, 97.5th, 99th] × [1st, 2.5th, 5th, 10th] percentiles), demonstrating robustness of forbidden zone detection to parameter choices.

**Table S4. Violator Enrichment.** Cell type enrichment analysis in forbidden zones. Columns: cell type, total count, violator count, expected count under null, fold enrichment, Fisher’s exact test p-value, FDR-corrected q-value.

**Table S5. Null Model Statistics.** Per-donor comparison of observed violation rates vs. permutation null expectation (n=1,000 permutations). Includes null mean, null SD, observed rate, z-score, enrichment ratio.

**Table S6. Violator QC.** Quality control comparison between violator and conformist cells. Columns: metric (ATAC counts, RNA counts, mitochondrial fraction, detected features), violator mean, conformist mean, Mann-Whitney p-value, effect size.

**Table S7. MI per Donor × State.** Empirical mutual information estimates by donor and developmental state, with 95% confidence intervals from bootstrap resampling (R=20). Includes cell counts per stratum.

**Table S8. MI vs Null (Permutation).** Comparison of empirical MI against blocked permutation null distributions (n=1,000 permutations per donor × state). Includes null mean, null SD, empirical MI, one-sided p-value.

**Table S9. Donor-Level Paired Inference.** ΔMI values (progenitor − differentiated) for each valid donor, directionality assessment, and Wilcoxon signed-rank test statistics. Documents the falsification of the informational dissociation hypothesis.

