## Supplementary material for "State-dependent geometric constraints reveal a regulatory gate in hematopoietic progenitors": data supplementary: README_reproducibility.md

# Reproducibility Manifest
**Date**: 2025-12-31T12:28:19.332027
**Status**: Frozen (NO-GO Verdict)

## Submission Package Content
This package contains the rigorous Falsification Report artifacts and the final figures/tables for the manuscript.

### Directory Structure
- `figures/`: Main manuscript figures (1-3) and source data.
- `tables/`: All supplementary tables (S1-S9).
- `supplementary/`: Supplementary MI analysis (PDF, Methods).
- `reports/`: The final Falsification Report with the NO-GO verdict.
- `scripts/`: Python scripts for reproduction.
- `metadata/`: File manifests and hashes.

### Scientific Verdict
The Mutual Information analysis (Table S7-S9) falsifies the hypothesis of informational decoupling in progenitors.
Findings indicate that geometric constraints (forbidden zones) co-occur with high informational coupling (efficiency).

### Reproduction
Run `python scripts/analysis_pipeline_mi.py` to reproduce MI calculations.
Run `python scripts/generate_mi_figure.py` to reproduce Supplementary Figure MI.

### Integrity
Verify file integrity using `metadata/run_hashes.sha256`:
`shasum -c metadata/run_hashes.sha256`
