## Supplementary material for "State-dependent geometric constraints reveal a regulatory gate in hematopoietic progenitors": data supplementary: Falsification_Report_MI.md

# Falsification Report: Chromatin-Transcription Decoupling

**Status**: [NO-GO]
**Date**: 2025-12-31

## 1. Executive Summary

This report presents a rigorous, "Science-proof" falsification analysis of the hypothesis that chromatin accessibility and gene expression are informatically decoupled in hematopoietic progenitors compared to solidified differentiated cells.

**Verdict**: **NO-GO (Falsified)**
**Key Finding**: Mutual Information (MI) analysis **does not support** a state-dependent reduction in coupling in progenitors. In fact, valid donors consistently show **higher** coupling in progenitors than in differentiated cells.
**Statistical Rigor**:
- Unit of Inference: **Donor (N=13 processed, N=5 valid)**
  - *Note: 8 donors excluded due to insufficient progenitor abundance (<500 cells) for robust inference.*
- Confounder Control: **Within-donor residualization**
- Sample Size: **Fixed N=2000 subsampling** (R=20 replicates)
- Null Model: **Blocked Permutation** (Donor, State)

---

## 2. Methodology Audit

| Component | Implementation | Status |
| :--- | :--- | :--- |
| **Inference Unit** | Donor-level (N=5 valid inference) | ✅ |
| **Confounder Control** | Residualized $A \sim QC$ and $T \sim QC$ *within* each donor | ✅ |
| **Estimator** | Copula Transform + KSG (k=5, 10, 20) | ✅ |
| **Sample Size** | Fixed $N=2000$ subsampling (R=20 reps) per state | ✅ |
| **Null Model** | Permuted $T$ within (Donor, State) blocks (n=1000) | ✅ |

---

## 3. Results

### 3.1 Empirical Mutual Information (Table S7)

*Data summarizes the 5 donors with sufficient progenitor cells.*

| Donor | $MI_{prog}$ (k=10) | $MI_{diff}$ (k=10) | $\Delta MI$ |
| :--- | :--- | :--- | :--- |
| **site2_donor4**| 0.0199 | 0.0183 | +0.0017 |
| **site3_donor10**| 0.0956 | 0.0109 | +0.0847 |
| **site4_donor1**| 0.0312 | 0.0227 | +0.0085 |
| **site4_donor8**| 0.0225 | 0.0192 | +0.0032 |
| **site4_donor9**| 0.0528 | 0.0163 | +0.0365 |

### 3.2 Falsification Test (Table S8)

Comparison against null distribution permuted within blocks.

**Progenitors (Aggregated):**
- Donors with $MI_{emp} > MI_{null}$ ($p < 0.05$): **4 / 5**
  - Significant coupling detected in 80% of valid donors.
- Mean $\Delta MI$ ($MI_{emp} - MI_{null}$): **Positive** (Range: +0.013 to +0.089)

**Differentiated (Aggregated):**
- Donors with $MI_{emp} > MI_{null}$ ($p < 0.05$): **5 / 5**
- Mean $\Delta MI$: **Positive** (Range: +0.004 to +0.012)

### 3.3 Donor-Level Paired Inference (Table S9)

Hypothesis: $\Delta MI = MI_{prog} - MI_{diff} < 0$.

- **N Donors**: 5 (with sufficient N)
- **Donors Corroborating ($\Delta < 0$)**: **0 / 5**
- **Median $\Delta MI$**: **+0.0085** (Progenitors have *higher* MI)
- **Wilcoxon Signed-Rank Test**: $p = 1.0$ (One-sided less)

---

## 4. Verdict and Recommendation

Based on the rigorous falsification criteria:

> **FAILED TO FALSIFY NULL HYPOTHESIS**
> The hypothesis that progenitors exhibit reduced informational coupling is **falsified**. The data supports the opposite: progenitors maintain higher chromatin-transcription coupling than differentiated cells, likely reflecting active developmental regulation. The "forbidden zones" observed geometrically do not correspond to entropic decoupling.

**Next Steps**:
- [ ] Move MI analysis to Supplementary Materials as a negative result / limitation.
- [ ] Refine the manuscript claim: "Geometric constraints exist (Forbidden Zones) but imply distinct regulatory modes rather than informational decoupling."
- [ ] Freeze `submission_package` with these findings.
