## Supplementary material for "State-dependent geometric constraints reveal a regulatory gate in hematopoietic progenitors": data supplementary: analysis_pipeline_mi.py

```
import os
import json
import numpy as np
import pandas as pd
import muon as mu
from scipy import stats
from sklearn.linear_model import LinearRegression
from sklearn.neighbors import NearestNeighbors
from scipy.special import digamma
import logging
import sys
from joblib import Parallel, delayed

# --- CONFIGURATION ---
BASE_DIR = r"c:\Users\ELKIN\.gemini\antigravity\scratch"
DATA_DIR = os.path.join(BASE_DIR, r"Permeability_Jamming\02_ingest\processed")
PROCESSED_JSON = os.path.join(BASE_DIR, r"Permeability_Jamming\02_ingest\processed_frozen.json")
OUTPUT_DIR = os.path.join(BASE_DIR, r"hematopoiesis_decoupling\results")
os.makedirs(OUTPUT_DIR, exist_ok=True)

# State Definitions
PROGENITOR_SET = {
    'HSC', 
    'Lymph prog', 
    'G/M prog', 
    'MK/E prog', 
    'Proerythroblast'
}

# --- LOGGING ---
# Logging in parallel is tricky. We'll rely on print which joblib captures or main process logging.
logging.basicConfig(
    filename=os.path.join(OUTPUT_DIR, "analysis_mi_science_parallel.log"),
    level=logging.INFO,
    format="%(asctime)s - %(levelname)s - %(message)s"
)
logger = logging.getLogger()
console = logging.StreamHandler(sys.stdout)
console.setLevel(logging.INFO)
logger.addHandler(console)

# --- UTILITIES ---

def load_dataset_metadata():
    with open(PROCESSED_JSON, 'r') as f:
        meta = json.load(f)
    return [m for m in meta if m['dataset_id'] == 'GSE194122']

def residualize_feature(y, X_reg):
    if X_reg is None or X_reg.shape[1] == 0:
        return y
    if np.isnan(X_reg).any():
        col_mean = np.nanmedian(X_reg, axis=0)
        inds = np.where(np.isnan(X_reg))
        X_reg[inds] = np.take(col_mean, inds[1])
    reg = LinearRegression().fit(X_reg, y)
    return y - reg.predict(X_reg)

def copula_transform(x, epsilon=1e-6):
    noise = np.random.uniform(-epsilon, epsilon, size=x.shape)
    x_jittered = x + noise
    ranks = stats.rankdata(x_jittered)
    return (ranks - 0.5) / len(ranks)

def ksg_mi(x, y, k=10):
    n = len(x)
    data = np.column_stack((x, y))
    nbrs = NearestNeighbors(n_neighbors=k + 1, algorithm='auto', metric='chebyshev').fit(data)
    distances, _ = nbrs.kneighbors(data)
    radius = np.maximum(distances[:, k], 1e-15) 
    
    from scipy.spatial import cKDTree
    tree_x = cKDTree(x.reshape(-1, 1))
    tree_y = cKDTree(y.reshape(-1, 1))
    
    nx = np.array([len(l) for l in tree_x.query_ball_point(x.reshape(-1, 1), radius - 1e-15, p=np.inf)])
    ny = np.array([len(l) for l in tree_y.query_ball_point(y.reshape(-1, 1), radius - 1e-15, p=np.inf)])
    
    psi_n = digamma(n)
    psi_k = digamma(k)
    psi_nx = np.mean(digamma(nx))
    psi_ny = np.mean(digamma(ny))
    
    return max(0.0, psi_k + psi_n - psi_nx - psi_ny)

# --- WORKER FUNCTION ---

def process_donor(meta_entry):
    donor_id = meta_entry['sample_id']
    fpath = os.path.join(DATA_DIR, meta_entry['file'])
    
    # Simple print for monitoring (joblib might clutter but it shows activity)
    print(f"Starting {donor_id}...")
    
    try:
        mdata = mu.read(fpath)
    except Exception as e:
        print(f"Failed {donor_id}: {e}")
        return []

    # 1. PREPARE
    if 'ATAC_atac_fragments' in mdata.mod['atac'].obs:
        A_raw = np.log1p(mdata.mod['atac'].obs['ATAC_atac_fragments'].values)
    else:
        A_raw = np.log1p(np.array(mdata.mod['atac'].X.sum(axis=1)).flatten())

    if 'GEX_n_counts' in mdata.mod['rna'].obs:
        T_raw = np.log1p(mdata.mod['rna'].obs['GEX_n_counts'].values)
    else:
        T_raw = np.log1p(np.array(mdata.mod['rna'].X.sum(axis=1)).flatten())
        
    if 'cell_type' in mdata.mod['rna'].obs:
        labels = mdata.mod['rna'].obs['cell_type'].values.astype(str)
    else:
        return []

    # 2. RESIDUALIZE
    atac_qc_cols = ['ATAC_atac_fragments', 'ATAC_reads_in_peaks_frac', 'ATAC_nucleosome_signal', 'ATAC_tss_enrichment']
    valid_atac = [c for c in atac_qc_cols if c in mdata.mod['atac'].obs]
    if valid_atac:
        X_atac = mdata.mod['atac'].obs[valid_atac].values
        A_res = residualize_feature(A_raw, X_atac)
    else:
        A_res = A_raw

    rna_qc_cols = ['GEX_n_counts', 'GEX_n_genes', 'GEX_pct_counts_mt']
    valid_rna = [c for c in rna_qc_cols if c in mdata.mod['rna'].obs]
    if valid_rna:
        X_rna = mdata.mod['rna'].obs[valid_rna].values
        T_res = residualize_feature(T_raw, X_rna)
    else:
        T_res = T_raw

    # 3. COPULA
    A_cop = copula_transform(A_res)
    T_cop = copula_transform(T_res)

    # 4. GROUPS
    groups = {}
    unique_states = np.unique(labels)
    for s in unique_states:
        groups[f"State_{s}"] = np.where(labels == s)[0]
    
    prog_mask = np.isin(labels, list(PROGENITOR_SET))
    groups["AGG_Progenitor"] = np.where(prog_mask)[0]
    groups["AGG_Differentiated"] = np.where(~prog_mask)[0]

    # 5. MI
    results = []
    for group_name, indices in groups.items():
        n_total = len(indices)
        if n_total < 500:
            continue
            
        n_fixed = 2000 if n_total >= 2000 else n_total
        
        reps_mi_k10 = []
        reps_mi_k5 = []
        reps_mi_k20 = []
        null_pool_values = []
        
        for r in range(20):
            if n_total > n_fixed:
                sub_idx = np.random.choice(indices, n_fixed, replace=False)
            else:
                sub_idx = indices 
            
            A_sub = A_cop[sub_idx]
            T_sub = T_cop[sub_idx]
            
            reps_mi_k10.append(ksg_mi(A_sub, T_sub, k=10))
            reps_mi_k5.append(ksg_mi(A_sub, T_sub, k=5))
            reps_mi_k20.append(ksg_mi(A_sub, T_sub, k=20))
            
            # Null (5 reps * 1000 perms)
            if r < 5:
                T_curr = T_sub.copy()
                # 1000 perms
                for _ in range(1000):
                    np.random.shuffle(T_curr)
                    # Inline calc for speed? No, use func.
                    # Optimization: KSG is slow. 
                    # T_curr is shuffled. A_sub fixed.
                    null_pool_values.append(ksg_mi(A_sub, T_curr, k=10))

        mi_emp_mean = np.mean(reps_mi_k10)
        n_greater = np.sum(np.array(null_pool_values) >= mi_emp_mean)
        p_perm = (1 + n_greater) / (1 + len(null_pool_values))
        
        results.append({
            'donor': donor_id,
            'state_label': group_name.replace("State_", "").replace("AGG_", ""),
            'is_aggregate': group_name.startswith("AGG_"),
            'n_total': n_total,
            'n_subsample': n_fixed,
            'mi_k10_mean': mi_emp_mean,
            'mi_k10_ci_low': np.percentile(reps_mi_k10, 5),
            'mi_k10_ci_high': np.percentile(reps_mi_k10, 95),
            'mi_k5_mean': np.mean(reps_mi_k5),
            'mi_k20_mean': np.mean(reps_mi_k20),
            'mi_null_mean': np.mean(null_pool_values),
            'mi_null_sd': np.std(null_pool_values),
            'p_perm': p_perm,
            'delta_mi': mi_emp_mean - np.mean(null_pool_values)
        })
    print(f"Finished {donor_id}")
    return results

def main():
    logger.info("Starting Parallel Science-Proof MI Analysis")
    metadata = load_dataset_metadata()
    logger.info(f"Loaded {len(metadata)} donors.")
    
    # Parallel Execution
    # n_jobs=6 to leave room for system
    all_results_nested = Parallel(n_jobs=6, verbose=10)(
        delayed(process_donor)(meta) for meta in metadata
    )
    
    # Flatten
    all_results = [item for sublist in all_results_nested for item in sublist]
            
    df = pd.DataFrame(all_results)
    out_path = os.path.join(OUTPUT_DIR, "mi_results_science_raw.tsv")
    df.to_csv(out_path, sep='\t', index=False)
    logger.info(f"Analysis complete. Results saved to {out_path}")

if __name__ == "__main__":
    main()
```
