## Supplementary material for "State-dependent geometric constraints reveal a regulatory gate in hematopoietic progenitors": data supplementary: freeze_package.py

```
import os
import hashlib
import json
import pandas as pd
from datetime import datetime

# --- CONFIGURATION ---
BASE_DIR = r"c:\Users\ELKIN\.gemini\antigravity\scratch\hematopoiesis_decoupling\submission_package"
METADATA_DIR = os.path.join(BASE_DIR, "metadata")
os.makedirs(METADATA_DIR, exist_ok=True)

def calculate_sha256(filepath):
    sha256_hash = hashlib.sha256()
    with open(filepath, "rb") as f:
        for byte_block in iter(lambda: f.read(4096), b""):
            sha256_hash.update(byte_block)
    return sha256_hash.hexdigest()

def main():
    print(f"Freezing Submission Package at: {BASE_DIR}")
    manifest_rows = []
    
    # Walk everything in submission_package
    for root, dirs, files in os.walk(BASE_DIR):
        if "metadata" in root: continue # Don't hash the manifest itself while creating it
        if "__pycache__" in root: continue
        
        for file in files:
            filepath = os.path.join(root, file)
            rel_path = os.path.relpath(filepath, BASE_DIR)
            
            size_bytes = os.path.getsize(filepath)
            sha256 = calculate_sha256(filepath)
            
            manifest_rows.append({
                'file_path': rel_path,
                'size_bytes': size_bytes,
                'sha256': sha256,
                'last_modified': datetime.fromtimestamp(os.path.getmtime(filepath)).isoformat()
            })
    
    # Write Manifest
    df = pd.DataFrame(manifest_rows)
    manifest_path = os.path.join(METADATA_DIR, "manifest_files.tsv")
    df.to_csv(manifest_path, sep='\t', index=False)
    print(f"Generated {manifest_path}")
    
    # Write Run Hashes (Simple Format)
    hashes_path = os.path.join(METADATA_DIR, "run_hashes.sha256")
    with open(hashes_path, 'w') as f:
        for _, row in df.iterrows():
            f.write(f"{row['sha256']}  {row['file_path']}\n")
    print(f"Generated {hashes_path}")
    
    # Write README_reproducibility
    readme_content = f"""# Reproducibility Manifest
**Date**: {datetime.now().isoformat()}
**Status**: Frozen (NO-GO Verdict)

## Submission Package Content
This package contains the rigorous Falsification Report artifacts and the final figures/tables for the manuscript.

### Directory Structure
- `figures/`: Main manuscript figures (1-3) and source data.
- `tables/`: All supplementary tables (S1-S9).
- `supplementary/`: Supplementary MI analysis (PDF, Methods).
- `reports/`: The final Falsification Report with the NO-GO verdict.
- `scripts/`: Python scripts for reproduction.
- `metadata/`: File manifests and hashes.

### Scientific Verdict
The Mutual Information analysis (Table S7-S9) falsifies the hypothesis of informational decoupling in progenitors.
Findings indicate that geometric constraints (forbidden zones) co-occur with high informational coupling (efficiency).

### Reproduction
Run `python scripts/analysis_pipeline_mi.py` to reproduce MI calculations.
Run `python scripts/generate_mi_figure.py` to reproduce Supplementary Figure MI.

### Integrity
Verify file integrity using `metadata/run_hashes.sha256`:
`shasum -c metadata/run_hashes.sha256`
"""
    readme_path = os.path.join(BASE_DIR, "README_reproducibility.md")
    with open(readme_path, 'w') as f:
        f.write(readme_content)
    print(f"Generated {readme_path}")

if __name__ == "__main__":
    main()
```
