## Supplementary material for "State-dependent geometric constraints reveal a regulatory gate in hematopoietic progenitors": data supplementary: generate_mi_figure.py

```
import pandas as pd
import matplotlib.pyplot as plt
import seaborn as sns
import os

# --- CONFIGURATION ---
BASE_DIR = r"c:\Users\ELKIN\.gemini\antigravity\scratch\hematopoiesis_decoupling"
OUTPUT_DIR = os.path.join(BASE_DIR, "submission_package/supplementary")
SOURCE_DATA = os.path.join(BASE_DIR, "figures/source_data/Fig_MI_State.tsv")

def main():
    if not os.path.exists(OUTPUT_DIR):
        os.makedirs(OUTPUT_DIR)
        
    print("Generating Supplementary Figure MI...")
    if not os.path.exists(SOURCE_DATA):
        print(f"Source data not found: {SOURCE_DATA}")
        return

    df = pd.read_csv(SOURCE_DATA, sep='\t')
    
    # Filter for Aggregates
    aggs = df[df['is_aggregate'] == True].copy()
    
    if aggs.empty:
        print("No aggregate data found.")
        return

    plt.figure(figsize=(6, 5))
    sns.boxplot(data=aggs, x='state_label', y='mi_k10_mean', palette='Set2')
    sns.stripplot(data=aggs, x='state_label', y='mi_k10_mean', color='black', alpha=0.7)
    
    plt.title("Mutual Information by State (Aggregated)")
    plt.ylabel("Mutual Information (k=10)")
    plt.xlabel("State")
    
    # Connect paired donors
    donors = aggs['donor'].unique()
    for d in donors:
        subset = aggs[aggs['donor'] == d]
        if len(subset) == 2:
            prog = subset[subset['state_label'] == 'Progenitor']['mi_k10_mean'].values
            diff = subset[subset['state_label'] == 'Differentiated']['mi_k10_mean'].values
            if len(prog) > 0 and len(diff) > 0:
                # Get x-coordinates
                # Progenitor is usually first or second depending on sort.
                # Let's assume alphabetical: Differentiated (0), Progenitor (1)
                # Check order matches plot
                # default sort is alpha
                y_diff = diff[0]
                y_prog = prog[0]
                plt.plot([0, 1], [y_diff, y_prog], color='gray', linestyle='--', alpha=0.5)

    plt.tight_layout()
    out_path = os.path.join(OUTPUT_DIR, "Supplementary_Fig_MI.pdf")
    plt.savefig(out_path)
    print(f"Saved {out_path}")

if __name__ == "__main__":
    main()
```
