## Supplementary material for "State-dependent geometric constraints reveal a regulatory gate in hematopoietic progenitors": data supplementary: generate_tables_mi.py

```
import pandas as pd
import numpy as np
import os
from scipy.stats import wilcoxon

# --- CONFIGURATION ---
RESULTS_DIR = r"c:\Users\ELKIN\.gemini\antigravity\scratch\hematopoiesis_decoupling\results"
OUTPUT_DIR = r"c:\Users\ELKIN\.gemini\antigravity\scratch\hematopoiesis_decoupling\submission_tables"
SOURCE_DATA_DIR = r"c:\Users\ELKIN\.gemini\antigravity\scratch\hematopoiesis_decoupling\figures\source_data"
os.makedirs(OUTPUT_DIR, exist_ok=True)
os.makedirs(SOURCE_DATA_DIR, exist_ok=True)

RAW_FILE = os.path.join(RESULTS_DIR, "mi_results_science_raw.tsv")

def load_data():
    if not os.path.exists(RAW_FILE):
        raise FileNotFoundError(f"{RAW_FILE} not found. Run analysis pipeline first.")
    return pd.read_csv(RAW_FILE, sep='\t')

def generate_table_s7(df):
    """
    Table S7: MI Statistics per Donor/State
    """
    # Filter detailed states + aggregates
    # Columns: Donor | State | N_Total | N_Sub | MI_Mean | MI_CI_Low | MI_CI_High | Sens_k5 | Sens_k20
    
    cols = ['donor', 'state_label', 'n_total', 'n_subsample', 'mi_k10_mean', 'mi_k10_ci_low', 'mi_k10_ci_high', 'mi_k5_mean', 'mi_k20_mean']
    t7 = df[cols].copy()
    
    # Format
    t7['MI (k=10, 95% CI)'] = t7.apply(lambda x: f"{x['mi_k10_mean']:.4f} [{x['mi_k10_ci_low']:.4f}-{x['mi_k10_ci_high']:.4f}]", axis=1)
    
    out_path = os.path.join(OUTPUT_DIR, "Table_S7_MI_Donor_State.tsv")
    t7.to_csv(out_path, sep='\t', index=False)
    print(f"Generated {out_path}")

def generate_table_s8(df):
    """
    Table S8: Null Hypothesis Stats
    """
    # Columns: Donor | State | MI_Emp | MI_Null_Mean | MI_Null_SD | P_Perm | Delta_MI
    cols = ['donor', 'state_label', 'mi_k10_mean', 'mi_null_mean', 'mi_null_sd', 'p_perm', 'delta_mi']
    t8 = df[cols].copy()
    
    t8.rename(columns={'mi_k10_mean': 'MI_Empirical'}, inplace=True)
    out_path = os.path.join(OUTPUT_DIR, "Table_S8_MI_Null_Stats.tsv")
    t8.to_csv(out_path, sep='\t', index=False)
    print(f"Generated {out_path}")

def generate_table_s9_and_verdict(df):
    """
    Table S9: Donor-Level Inference (Paired Test)
    Computes Wilcoxon Signed-Rank Test on Delta MI (Prog - Diff).
    """
    # Pivot to get Prog and Diff for each donor
    # We use the AGGREGATE labels: 'Progenitor' and 'Differentiated'
    # Note in pipeline: label is result of strip("AGG_") so "Progenitor", "Differentiated"
    
    # Filter for aggregates
    aggs = df[df['is_aggregate'] == True]
    
    piv = aggs.pivot(index='donor', columns='state_label', values='mi_k10_mean')
    
    if 'Progenitor' not in piv.columns or 'Differentiated' not in piv.columns:
        print("Warning: Missing Progenitor or Differentiated columns in aggregates.")
        return
    
    res = piv[['Progenitor', 'Differentiated']].dropna()
    res['Delta_MI'] = res['Progenitor'] - res['Differentiated']
    res['Verdict_Local'] = res['Delta_MI'] < 0
    
    # Test
    stats_res = wilcoxon(res['Progenitor'], res['Differentiated'], alternative='less')
    p_val = stats_res.pvalue
    n_donors = len(res)
    n_corroborated = res['Verdict_Local'].sum()
    median_delta = res['Delta_MI'].median()
    
    # Save Table S9
    res.reset_index(inplace=True)
    out_path = os.path.join(OUTPUT_DIR, "Table_S9_Donor_Level_Inference.tsv")
    res.to_csv(out_path, sep='\t', index=False)
    print(f"Generated {out_path}")
    
    # Print Verdict Info to stdout (for capture)
    print("\n--- VERDICT DATA ---")
    print(f"N Donors: {n_donors}")
    print(f"Corroborated (Delta < 0): {n_corroborated}")
    print(f"Median Delta: {median_delta:.5f}")
    print(f"Wilcoxon p-value: {p_val:.5e}")
    if n_corroborated >= 9 and p_val < 0.05:
        print("VERDICT: GO (Corroborated)")
    else:
        print("VERDICT: NO-GO (Falsified/Weak)")
    print("--------------------\n")

def generate_source_data(df):
    """
    Source Data for Figures
    """
    # Fig_MI_State.tsv -> same as raw basically but clean
    out1 = os.path.join(SOURCE_DATA_DIR, "Fig_MI_State.tsv")
    df.to_csv(out1, sep='\t', index=False)
    
    # Fig_MI_Null_vs_Emp.tsv -> S8 content basically
    cols = ['donor', 'state_label', 'mi_k10_mean', 'mi_null_mean', 'mi_null_sd']
    out2 = os.path.join(SOURCE_DATA_DIR, "Fig_MI_Null_vs_Emp.tsv")
    df[cols].to_csv(out2, sep='\t', index=False)
    print("Generated Source Data.")

def main():
    try:
        df = load_data()
        generate_table_s7(df)
        generate_table_s8(df)
        generate_table_s9_and_verdict(df)
        generate_source_data(df)
    except Exception as e:
        print(f"Error: {e}")

if __name__ == "__main__":
    main()
```
