## Supplementary material for "State-dependent geometric constraints reveal a regulatory gate in hematopoietic progenitors": data supplementary: package_assembler.py

```
import os
import shutil
import sys

# --- CONFIGURATION ---
BASE_DIR = r"c:\Users\ELKIN\.gemini\antigravity\scratch\hematopoiesis_decoupling"
TARGET_DIR = os.path.join(BASE_DIR, "submission_package")
SOURCE_PERMEABILITY = r"c:\Users\ELKIN\.gemini\antigravity\scratch\Permeability_Jamming\submission_package_FINAL"
SOURCE_TABLES_MI = os.path.join(BASE_DIR, "submission_tables")
SOURCE_REPORTS = os.path.join(BASE_DIR, "..", "..", "brain", "35135939-638c-42e2-bf91-676520b416d2") # Falsification report is an artifact
SOURCE_SCRIPTS = os.path.join(BASE_DIR, "scripts")
SOURCE_DATA_MI = os.path.join(BASE_DIR, "figures", "source_data")

# --- STRUCTURE ---
STRUCTURE = [
    "figures",
    "figures/source_data",
    "tables",
    "supplementary",
    "reports",
    "scripts",
    "scripts/utils",
    "metadata"
]

def clean_and_create():
    if os.path.exists(TARGET_DIR):
        print(f"Cleaning {TARGET_DIR}...")
        shutil.rmtree(TARGET_DIR)
    
    for d in STRUCTURE:
        os.makedirs(os.path.join(TARGET_DIR, d), exist_ok=True)
        
def copy_file(src, dst_dir, dst_name=None):
    if not os.path.exists(src):
        print(f"Error: Source not found {src}")
        return
    
    filename = os.path.basename(src) if dst_name is None else dst_name
    dst = os.path.join(dst_dir, filename)
    shutil.copy2(src, dst)
    print(f"Copied {filename} -> {dst_dir}")

def main():
    print("Assembling Submission Package...")
    clean_and_create()
    
    # 1. FIGURES (1-3)
    print("\n--- Migrating Figures 1-3 ---")
    fig_src = os.path.join(SOURCE_PERMEABILITY, "figures")
    for f in ["Figure_1_Validity.pdf", "Figure_2_The_Law.pdf", "Figure_3_The_Gate.pdf"]:
        copy_file(os.path.join(fig_src, f), os.path.join(TARGET_DIR, "figures"))
        
    # Source Data 1-3
    sd_src = os.path.join(fig_src, "source_data")
    if os.path.exists(sd_src):
         for f in os.listdir(sd_src):
             copy_file(os.path.join(sd_src, f), os.path.join(TARGET_DIR, "figures/source_data"))
    
    # 2. TABLES
    print("\n--- Migrating Tables ---")
    # S1-S6
    tab_src_perm = os.path.join(SOURCE_PERMEABILITY, "tables")
    for i in range(1, 7):
        copy_file(os.path.join(tab_src_perm, f"Table_S{i}_*.tsv"), os.path.join(TARGET_DIR, "tables")) # Pattern match needed?
        # Actually I saw the filenames, they are predictable.
        # But wait, globbing is safer.
        import glob
        files = glob.glob(os.path.join(tab_src_perm, f"Table_S{i}_*.tsv"))
        for f in files:
            copy_file(f, os.path.join(TARGET_DIR, "tables"))

    # S7-S9 (MI)
    for f in os.listdir(SOURCE_TABLES_MI):
        if f.endswith(".tsv"):
            copy_file(os.path.join(SOURCE_TABLES_MI, f), os.path.join(TARGET_DIR, "tables"))

    # 3. SUPPLEMENTARY
    print("\n--- Migrating Supplementary ---")
    # Figures/Source Data for MI
    if os.path.exists(SOURCE_DATA_MI):
        for f in os.listdir(SOURCE_DATA_MI):
            copy_file(os.path.join(SOURCE_DATA_MI, f), os.path.join(TARGET_DIR, "figures/source_data"))

    # 4. REPORTS
    print("\n--- Migrating Reports ---")
    # Falsification Report
    report_src = os.path.join(SOURCE_REPORTS, "falsification_report_MI.md")
    if os.path.exists(report_src):
        copy_file(report_src, os.path.join(TARGET_DIR, "reports"), "Falsification_Report_MI.md")
    else:
        print("Warning: Falsification Report artifact not found in brain.")

    # 5. SCRIPTS
    print("\n--- Migrating Scripts ---")
    scripts_to_copy = [
        "analysis_pipeline_mi.py",
        "generate_tables_mi.py",
        "freeze_package.py",
        "package_assembler.py", # Self
        "generate_mi_figure.py" # Planned
    ]
    for s in scripts_to_copy:
        src = os.path.join(SOURCE_SCRIPTS, s)
        if os.path.exists(src):
            copy_file(src, os.path.join(TARGET_DIR, "scripts"))
            
    # 6. README
    # Will be generated by freeze_package.py in the final step, but we can copy the temp one if needed.
    
    print("\nPackage Assembled.")

if __name__ == "__main__":
    main()
```
