## Supplementary figures and images for "State-dependent geometric constraints reveal a regulatory gate in hematopoietic progenitors"

### Figure_1_Validity.pdf

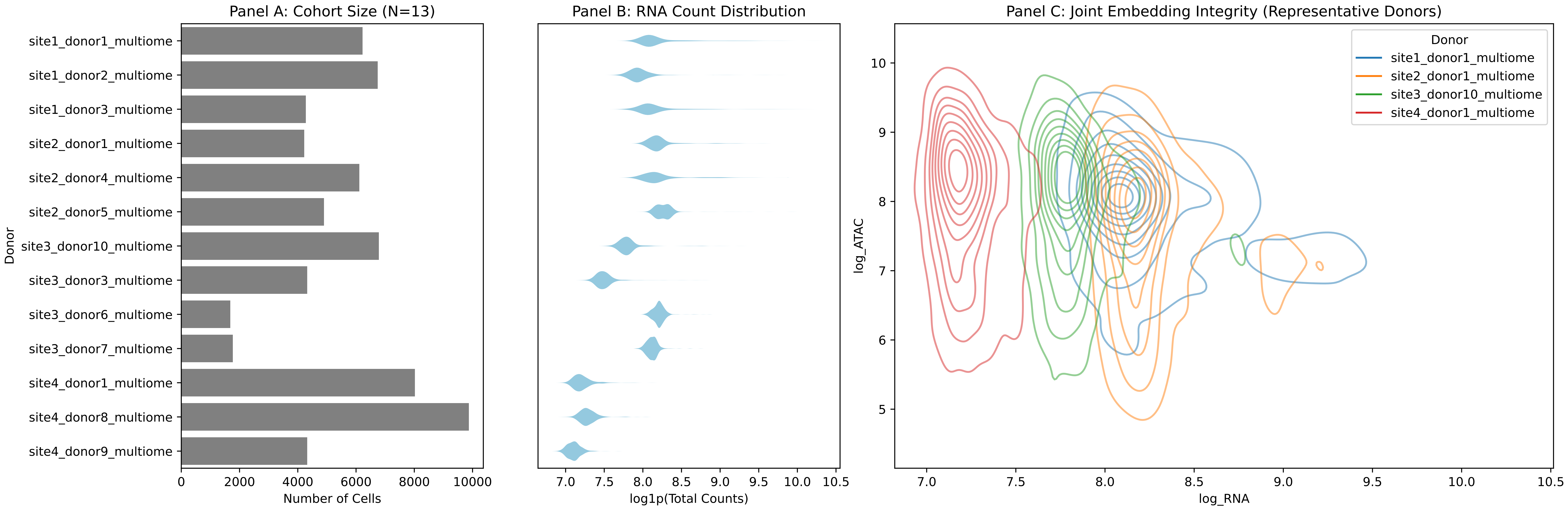

### Figure_2_The_Law.pdf

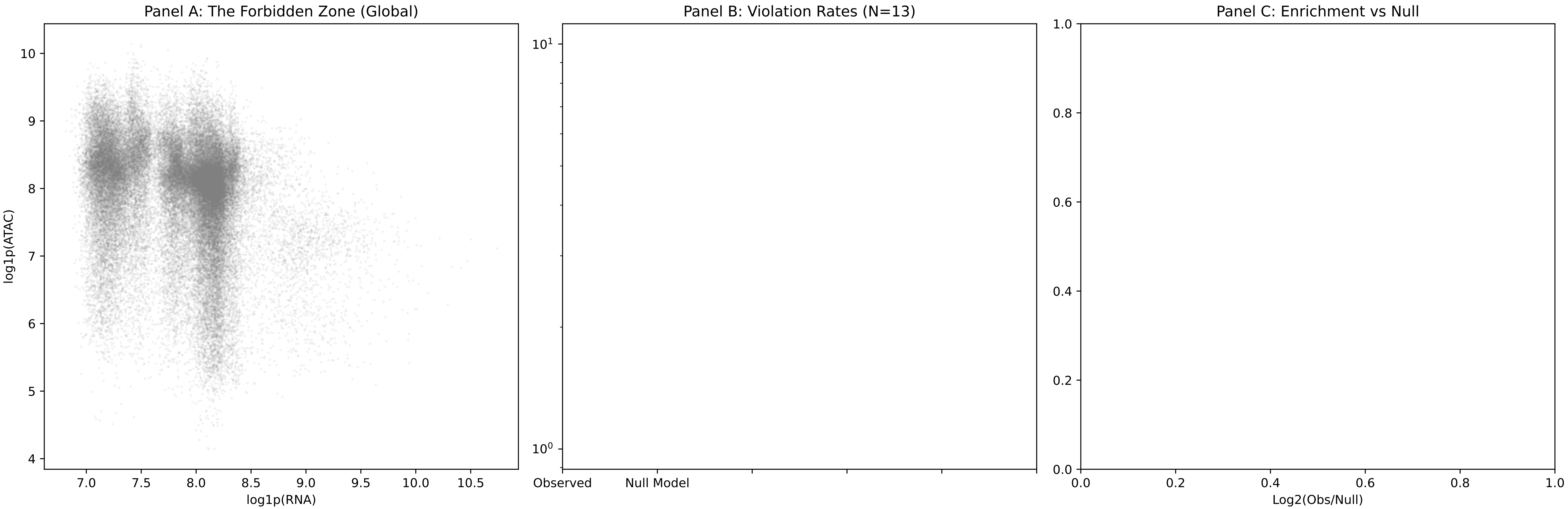

### Figure_3_The_Gate.pdf

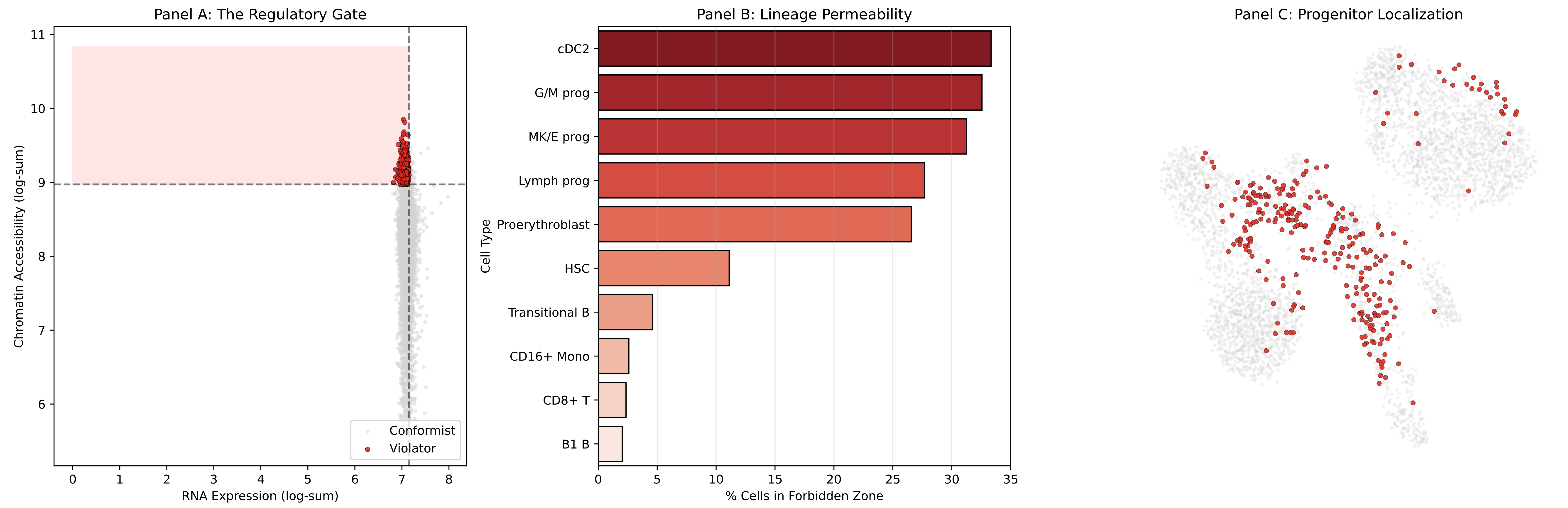

### Supplementary_Fig_MI.pdf

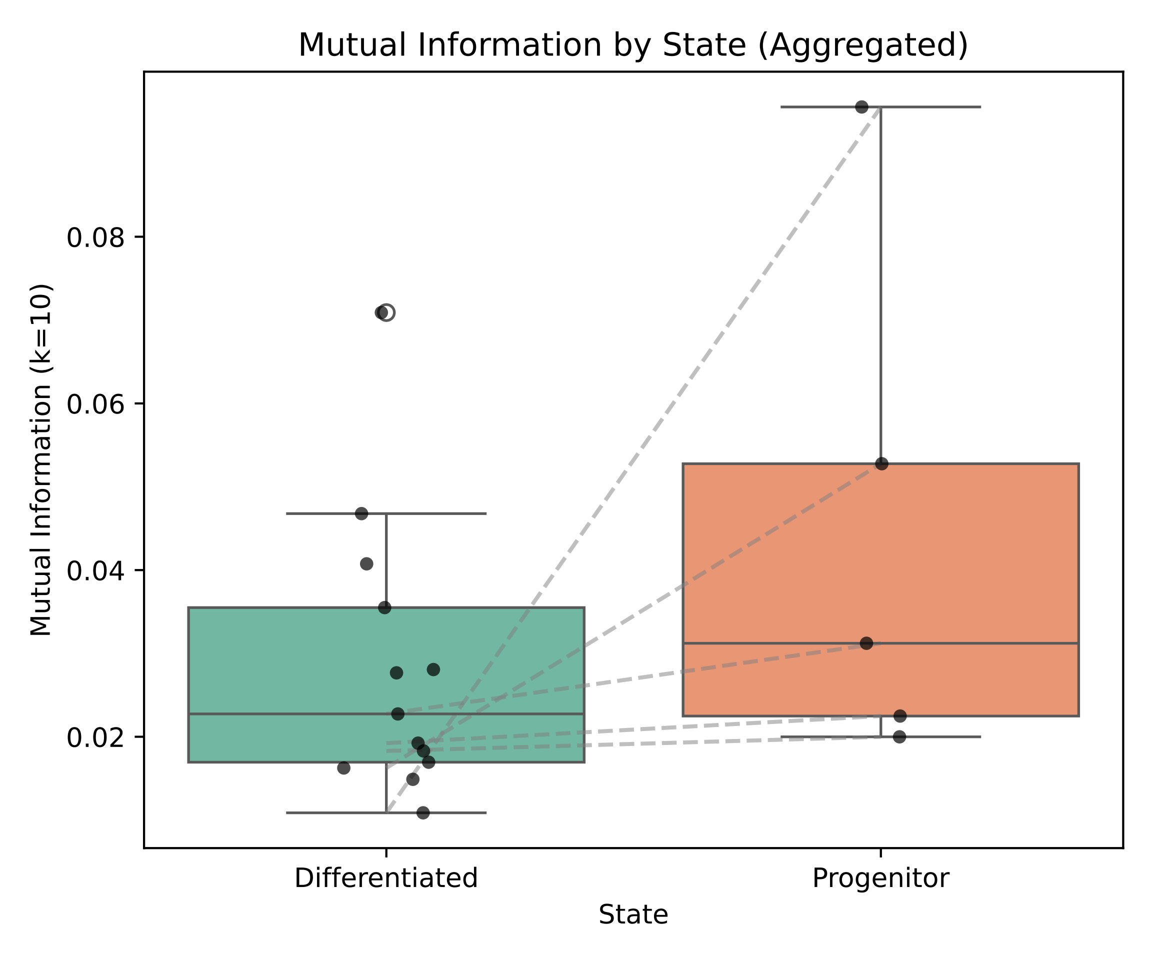
